# Decoding task-specific cognitive states with slow, directed functional networks in the human brain

**DOI:** 10.1101/681544

**Authors:** Devarajan Sridharan, Shagun Ajmera, Hritik Jain, Mali Sundaresan

**Affiliations:** Centre for Neuroscience, Indian Institute of Science, Bangalore 560012, India; Department of Computer Science and Automation, Indian Institute of Science, Bangalore 560012, India

**Keywords:** functional connectivity, Granger causality, machine learning, support vector machines, fMRI decoding, emergent dynamics, cognitive score prediction

## Abstract

Flexible functional interactions among brain regions mediate critical cognitive functions. Such interactions can be measured from functional magnetic resonance imaging (fMRI) data with either instantaneous (zero-lag) or lag-based (time-lagged) functional connectivity; only the latter approach permits inferring directed functional interactions. Yet, the fMRI hemodynamic response is slow, and sampled at a timescale (seconds) several orders of magnitude slower than the underlying neural dynamics (milliseconds). It is, therefore, widely held that lag-based fMRI functional connectivity, measured with approaches like as Granger-Geweke causality (GC), provides spurious and unreliable estimates of underlying neural interactions. Experimental verification of this claim has proven challenging because neural ground truth connectivity is often unavailable concurrently with fMRI recordings. We address this challenge by combining machine learning with GC functional connectivity estimation. We estimated instantaneous and lag-based GC functional connectivity networks using fMRI data from 1000 participants, drawn from the Human Connectome Project database. A linear classifier, trained on either instantaneous or lag-based GC, reliably discriminated among seven different task and resting brain states, with over 80% cross-validation accuracy. With network simulations, we demonstrate that instantaneous and lag-based GC exploited interactions at fast and slow timescales, respectively, to achieve robust classification. With human fMRI data, instantaneous and lag-based GC identified distinct, cognitive core networks. Finally, variations in GC connectivity explained inter-individual variations in a variety of cognitive scores. Our findings show that instantaneous and lag-based methods reveal complementary aspects of functional connectivity in the brain, and suggest that slow, directed functional interactions, estimated with fMRI, provide robust markers of behaviorally relevant cognitive states.

**Author Summary:** Functional MRI (fMRI) is a leading, non-invasive technique for mapping networks in the human brain. Yet, fMRI signals are noisy and sluggish, and fMRI scans are acquired at a timescale of seconds, considerably slower than the timescale of neural spiking (milliseconds). Can fMRI, then, be used to infer dynamic processes in the brain such as the direction of information flow among brain networks? We sought to answer this question by applying machine learning to fMRI scans acquired from 1000 participants in the Human Connectome Project (HCP) database. We show that directed brain networks, estimated with a technique known as Granger-Geweke Causality (GC), accurately predicts individual subjects’ task-specific cognitive states inside the scanner, and also explains variations in a variety of behavioral scores across individuals. We propose that directed functional connectivity, as estimated with fMRI-GC, is relevant for understanding human cognitive function.

## Introduction

Coordinated activity among brain regions underlies a variety of cognitive functions [1]. Mapping functional coupling among brain regions is, therefore, key to mapping brain function and for understanding how the brain produces behavior. Human fMRI studies have commonly investigated such functional coupling with correlation-based measures, including the Pearson correlation coefficient [2,3] and partial correlations [4] between pairs of brain regions. These measures frequently incorporate regularization penalties to estimate sparse functional networks [5]. Correlation-based measures are ideal for characterizing “instantaneous” functional coupling, representing functional interactions among brain regions that occur at timescales faster than the sampling rate of the measurement [6]. In contrast, comparatively few studies, have examined lag-based measures of functional connectivity to examine time-lagged interactions among brain regions [7,8].

Measures of linear dependence and feedback, based on Granger-Geweke causality (GC) [9,10] represent a powerful approach for estimating both instantaneous and lag-based functional connectivity. These measures are firmly grounded in information theory and statistical inferential frameworks [9–11]. GC measures have been widely applied to estimate functional connectivity in recordings of brain activity made with electroencephalography (EEG; [12]), magnetoencephalography (MEG; [13]) and electrocorticography (ECoG; [14]). However, the application of GC measures to brain recordings made with functional magnetic resonance imaging (fMRI) has provoked significant controversy [15–18]. Because the hemodynamic response is produced and sampled at a timescale (seconds) several orders of magnitude slower than the underlying neural processes (milliseconds), previous studies have argued that lag-based measures, particularly lag-based GC, produce spurious and unreliable estimates of functional connectivity, when applied to fMRI data (fMRI-GC) [18–21].

Three primary confounds have been identified with inferring connectivity with fMRI-GC. First, systematic differences in hemodynamic lags across regions could yield spurious directionality of GC connections [16,18]. Second, in simulations, measurement noise added to the signal during fMRI acquisition significantly degrades GC functional connectivity estimates [19]. Finally, downsampling recordings to the typical fMRI sampling rate (seconds), three orders of magnitude slower than the timescale of neural spiking (milliseconds), effectively eliminates all traces of functional connectivity inferred by GC [19].

The controversy regarding the application of GC to fMRI data continues to date, primarily because of the lack of access to ground-truth in neural data. On the one hand, claims regarding the efficacy of GC estimates are primarily based on simulations [11,22], and are only as valid as the underlying model of neural activity and hemodynamic responses. Because the precise mechanism by which neural responses generate hemodynamic responses is an active area of research, strong conclusions cannot be drawn based on fMRI simulations alone. On the other hand, establishing ground-truth validity for fMRI functional connectivity requires invasive neurophysiological recordings across many brain regions, concurrently during fMRI scans, a prohibitive enterprise.

We seek to address this controversy by applying machine learning to fMRI-GC networks, which works around these challenges. We estimated instantaneous and lag-based GC connectivity with fMRI data drawn from 1000 human subjects, recorded under seven different task conditions and in the resting state (∼8000 functional scans drawn from the Human Connectome Project database; [23]). We trained a linear classifier, based on GC connectivity features alone, to discriminate among the different task and resting conditions, and assessed classifier accuracy with cross validation. The results show that instantaneous and lag-based GC connectivity, estimated from fMRI data, can decode task-specific cognitive states with superlative accuracies. Next, with simulations, we show that slow, multi-second timescales emerge in sparse, random networks despite individual neurons operating at fast, millisecond timescales – a result that explains why directed functional connectivity can be reliably estimated with GC in slowly sampled fMRI data. Finally, we demonstrate that GC connectivity features can be used as predictors [24] to explain inter-individual variations in behavioral scores across a variety of cognitive tests. The results suggest that instantaneous and lag-based GC measures applied to fMRI data permit mapping slow, emergent and behaviorally relevant functional interactions in the human brain.

## Results

### GC estimated from slowly sampled fMRI data suffices to distinguish task and resting states

We asked if instantaneous GC (iGC) and directed GC (dGC) (SI Mathematical Note Section S1) connectivity would flexibly reconfigure with task demand, by testing if GC connectivity sufficed to accurately classify among seven different task states or the resting state (SI Table S1A; Methods; [9,10]). Data were obtained from 1000 participants from the Human Connectome Project (HCP) database [25] (RRID: SCR_008749). We used connection weights among brain regions in each network (iGC or dGC) as feature vectors in a linear classifier based on Support Vector Machines (SVM) for high dimensional predictor data. Accuracies for classifying resting state from a working memory task (WM task) are described first; accuracies for other tasks are presented subsequently.

Both iGC and dGC connectivity were able to distinguish the working memory task from resting state significantly above chance (Fig. 1B; p<0.001, permutation test). Maximum accuracy (median, [95% CI]) was 97.3% [96.3 - 98.0%] with iGC and 92.0% [90.5 - 93.2%] with dGC (SI Fig. S1B Yeo Parcellation, iGC: precision= 97.2, recall= 97.4; dGC: precision= 90.9, recall= 93.2). k-fold (k=10) cross-validation accuracy was comparable (iGC: 97.1% [96.2 - 97.9%], dGC: 91.7% [90.3 - 93.0%]). These numbers correspond to maximum cross validation accuracy across all five parcellations tested (SI Table S3; SI Fig. S1A); accuracies with each parcellation are shown in the Supporting Information (SI Fig. S1B).Non-linear classifiers, such as SVMs based on radial basis function kernels produced similar results, with comparably above chance classification accuracy for both iGC and dGC connectivity (SI Fig. S1C).

**Figure 1.**
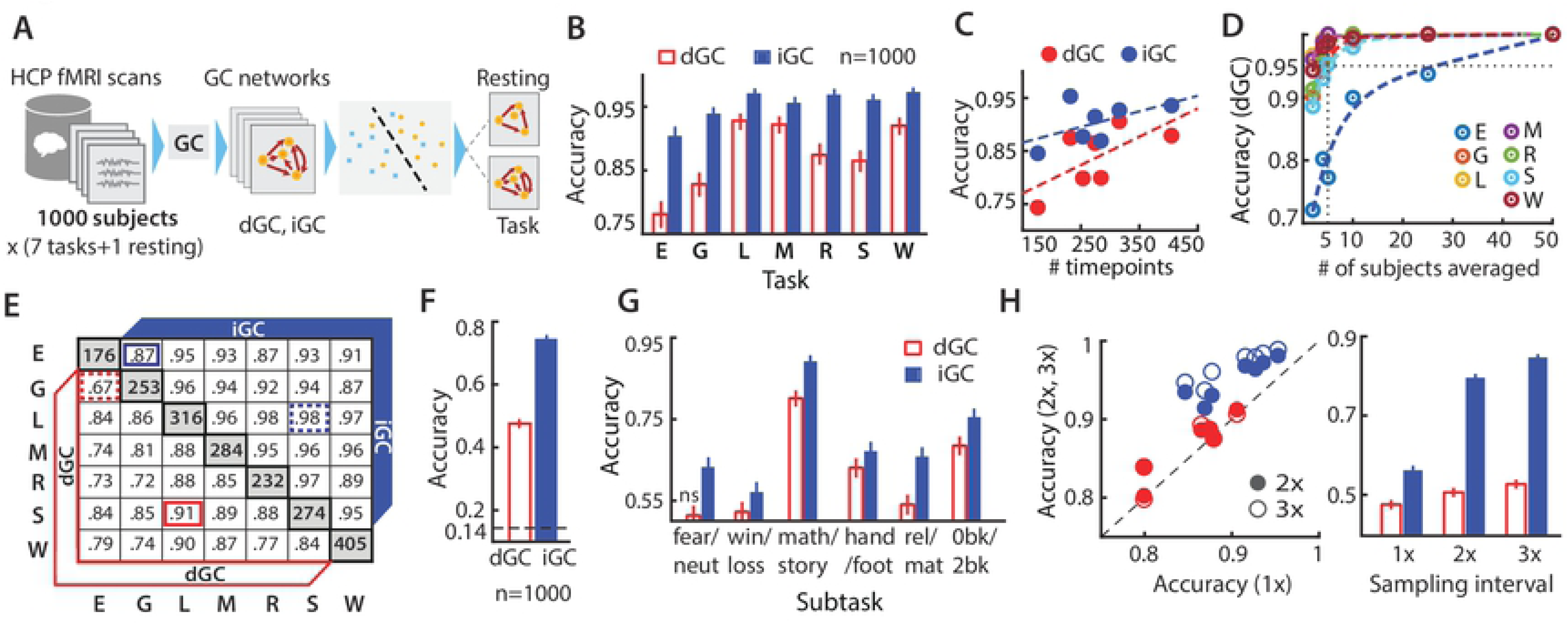
Discriminating between task and resting state with instantaneous and directed GC networks. **A.** Schematic of task state classification based on instantaneous (iGC) and directed (dGC) Granger-Geweke Causality with fMRI data from 1000 subjects (see text for details). **B.** Two-way classification accuracies (leave-one-out) for each of seven tasks versus resting state based on GC. Red unfilled bars and blue filled bars: accuracies based on dGC and iGC features, respectively (task key in SI Table S1). Error-bars: Clopper-Pearson binomial confidence intervals. Chance accuracy: 0.5 (not shown). **C.** Two-way task versus resting state classification accuracies based on dGC (red dots) and iGC (blue - dots), as a function of number of task scan time points (volumes). Dashed lines: linear fits. **D.** Two-way task versus resting state classification accuracies based on dGC after averaging dGC matrices over different numbers of subjects (x-axis). Each task is represented with a different color. Colored dashed lines: biexponential fits. Black dashed horizontal and vertical lines: 95% accuracy and n=5 subjects’ average, respectively. **E.** Two-way classification accuracies across each pair of tasks. Cells: classification accuracies for each pair of tasks based on dGC (lower triangular matrix) or iGC (upper triangular matrix). Diagonal cells: number of task scan timepoints. Highlighted cells: lowest (dashed-line border) and highest (solid-line border) accuracies achieved with dGC (red) and iGC (blue). **F.** N-way classification accuracies among all seven tasks. Dashed line: chance accuracy (14.3%). Other conventions are the same as in panel B. **G.** Two-way sub-task classification accuracies (SI Table S1B) based on GC. ns.: accuracy not significantly above chance. Other conventions are the same as in panel B. **H.** (Left) Two-way task versus resting state classification accuracies obtained with regional time series sub-sampled at 2x (filled symbols) and 3x (open symbols) of the TR (720 ms) (y-axis) plotted against accuracies obtained with the original data (1x, x-axis) for each of 7 tasks. Red: dGC, Blue: iGC. Dashed diagonal line: Line of equality (x=y). (Right) N-way classification accuracies among all seven tasks with data sampled at 1x, 2x, 3x of the original TR. Other conventions are the same as in panel F.For panels B,E,F: accuracies correspond to highest values, across all parcellations tested, and hyperparameter optimization was done for panel B. For panels C,G,H: accuracies correspond to Shirer et al [26] 14-network parcellation. For panel D: accuracies correspond to Shirer et al [26] 90-node parcellation.

We repeated these analyses by classifying the six other tasks (SI Table S1) versus resting state. iGC and dGC connectivity could accurately classify each task from resting state significantly above chance. For iGC, maximum classification accuracies ranged from 90.1%, for emotion task vs. resting state classification, to 97.1%, for language task vs. resting state classification. Similarly, for dGC, accuracies ranged from 78.1%, for emotion task vs. resting state classification, to 92.8%, for language task vs. resting state classification (Fig. 1B). In general, classification accuracy increased with more scan timepoints for each task versus resting state classification (Fig. 1C), consistent with GC being an information theoretic measure; we confirmed this result with simulations also (SI Fig. S1D).

In these analyses, classification accuracies based on dGC were systematically lower than those based on iGC. We asked if dGC accuracies were poorer due to noise corrupting the fit of the autoregressive model, and if a more consistent estimate could be obtained by averaging dGC connectivity features, to remove uncorrelated noise, across subjects. We addressed this question by partitioning the data into two groups -- a training (T) group and a test (S) groups – with 500 subjects each. We trained the classifier on group T and tested the classifier prediction by averaging GC matrices across several folds of S, each fold containing a few (m=2, 4, 5, 10, 25 or 50) subjects; the procedure was repeated by exchanging training and test datasets (see Methods). For the vast majority of tasks (6/7), dGC’s classification accuracy was more than 95% with as few as m=5 subjects within each fold of the test set (Fig. 1D). These results suggest that averaging dGC matrices across a few subjects, yielded reliable estimates of dGC connectivity.

We considered other factors that, in addition to intrinsic connectivity differences, could have produced these superior classification accuracies. First, GC-based accuracies for classifying task versus resting state scans might arise from differences in brain regions activated during each of these scans. In addition to task-relevant sensory input, overt motor responses always occurred during task scans but were absent during resting state scans [23,27]. Could GC features discriminate among more subtle connectivity variations across/within tasks? Second, scan data from the HCP database was sampled at a TR (repetition time) of 720 ms, considerably faster than the TR for conventional fMRI scans. Would GC accuracies degrade if the data were sampled at much slower sampling rate (∼2000 ms), in line with conventional fMRI TR?

We addressed the first question in two stages. First, we asked if GC connectivity features would be able to classify which of the seven tasks each subject was performing in the scanner. First, we performed a pairwise classification of each task from the other. Maximum classification accuracies for iGC (dGC) ranged from 87% (67%) for the emotion vs. gambling task classification to 98% (91%) for the language vs. social task classification. Again, the number of timepoints for each task proved to be a strong indicator of classification accuracies (Fig. 1E): average inter-task classification accuracies were highest for the language task (iGC: 97%, dGC:88%, n=316 timepoints) and lowest for the emotion task (iGC: 91%, dGC: 77%, n=176 timepoints). Next, we performed an n-way classification analysis across all 7 tasks, again using linear SVM (Methods). Accuracies were significantly above chance (14.3% for 1-in-7 classification) for classifying among the seven tasks (Fig. 1F; maximum accuracy, iGC: 74.4% [73.3%-75.4%]; dGC: 47.6% [46.4%-48.7%]; p<0.001, permutation test). These results indicate that functional connectivity was consistently estimated with GC, and reliably different across tasks.

Second, each of the different tasks in the HCP database comprised of blocks of contiguous trials, each corresponding to one of (at least) two different sub-tasks ([27]; SI Table S1B). For example, the motor task comprised of blocks of movements of the right or left hand interleaved with blocks of trials involving movement of the right or left foot. Similarly, the working memory task comprised of interleaved blocks of 0- back and 2-back tasks. We asked, therefore, if GC connectivity could distinguish among subtler variations in brain states across sub-tasks within each task. We sought to classify across two sub-tasks for each of six tasks (SI Table S1B). In all cases, except one, both iGC and dGC connectivity discriminated between each pair of sub-tasks with higher than chance accuracies (Fig. 1G; maximum accuracy, iGC: 89.2% [87.6% - 90.7%]; dGC: 80.1% [78.9% - 82.9%]; p<0.05 permutation test). These results indicate that GC functional connectivity could accurately distinguish among sub-tasks within each task as well.

Next, we tested whether GC connectivity estimated from slowly sampled fMRI data could accurately classify task and resting states. We downsampled the data to either one half (2xTR=1440 ms) or one third (3x TR=2160 ms) of its original sampling rate, by decimation, while also concatenating the decimated data to the end of the sub-sampled timeseries to preserve the overall number of timepoints (Methods). We repeated both of the previous classification analyses – pairwise task versus resting state classification (Fig. 1H left), as well as n-way inter-task classification (Fig. 1H right). Following downsampling, we observed that classification accuracies were marginally higher than accuracies in the original data in the case of dGC (2x: p=0.02; 3x: p=0.06; Wilcoxon one-tailed signed rank test), and were even higher than those in the original data in the case of iGC (2x: p=0.01; 3x: p=0.01), across tasks. These results indicate that the superlative sampling rate of the HCP fMRI data was not the primary reason for these high classification accuracies for GC-based classification.

We performed three control analyses to further confirm these results: i) by performing stationarity tests on the data prior to GC estimation and classification; ii) using single full regression to estimate GC [28,29], instead of estimating with separate full and reduced regressions; and iii) incorporating motion scrubbing [30] to ensure that the classification accuracies were not driven by head motion artifacts. These controls are described in SI Results, section on “Control analyses”. In every case, we obtained equivalent or superior classification accuracies (SI Fig. S2), confirming that the results were not due to data non-stationarity, biases in GC estimation or head motion artifacts.

These results demonstrate that both iGC and dGC yielded task-specific signatures of functional connectivity even with slowly sampled fMRI data (TR∼2000 ms): these estimates were consistent across subjects and reliably different across tasks to permit successful classification. Furthermore, these superlative classification accuracies were obtained despite widely held caveats concerning the application of GC to fMRI data [28].

### Correlation-purged GC connectivity suffices for accurate task-state classification

Correlation-based (zero-lag) connectivity measures (e.g. partial correlations or PC) have been widely applied to estimate functional connectivity from fMRI data [5,31]. In fact, several previous studies [18,19] have argued that correlation-based measures are more reliable and should be preferred to lag-based measures like GC [11], for estimating functional connectivity with fMRI data. We tested this claim here with a three-fold analysis approach.

First, we asked how classification accuracies based on PC connectivity would compare with those reported above, based on GC connectivity. Maximum classification accuracies with PC connectivity ranged from 96-99% for task versus resting state classification, and were consistently higher than accuracies with GC connectivity (Fig. 2A). These results are along expected lines: estimators based on same-time covariance, such as PC, are less susceptible to noise than those based on lagged covariance, such as GC (derived analytically in the Methods, section on *Functional connectivity estimation and classification with partial correlations*). In addition, as mentioned previously, GC is an information theoretic measure: classification accuracy with iGC and dGC increased systematically with more scan time points, asymptotically matching PC accuracies (SI Fig. S1D).

**Figure 2.**
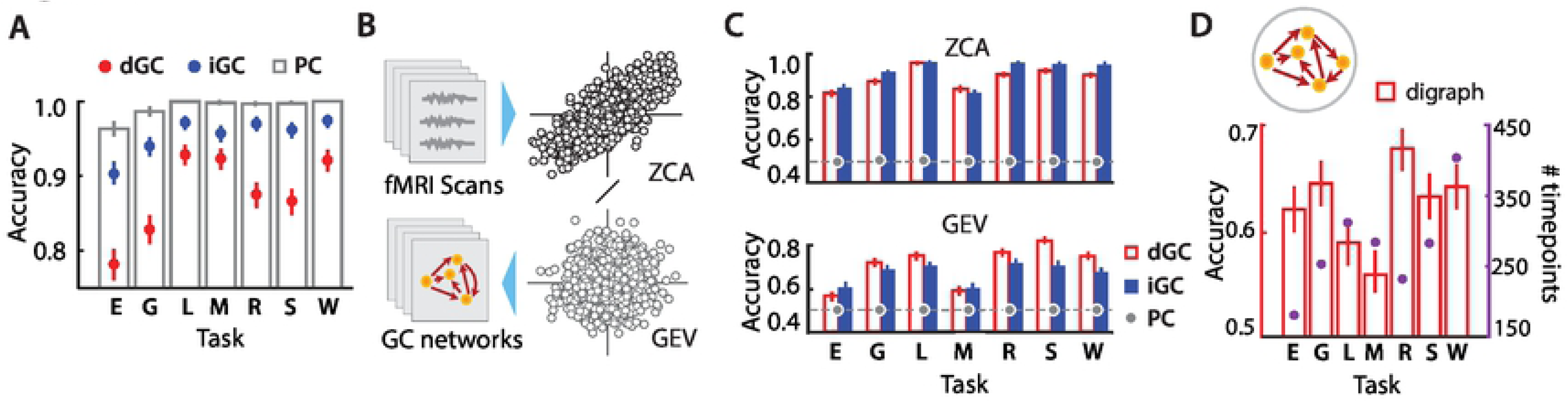
Classification accuracies with GC purged of instantaneous correlations. **A.** Two-way task versus resting state classification accuracies, based on partial correlations (PC; grey unfilled bars). Numbers reported correspond to highest leave-one-out classification accuracies across parcellations, obtained with hyperparameter optimization. Corresponding accuracies for dGC (red dots) and iGC (blue dots) are shown for comparison. Other conventions are as in Fig. 1B. **B.** Schematic illustrating procedure for purging data of instantaneous correlations. fMRI regional timeseries were purged of instantaneous correlations by either whitening the data with zero-phase component analysis (ZCA), separately for each task and resting state scan, or by projecting data into a space spanned by the generalized eigenvectors (GEV), common to both task and resting state scans. GC and PC were then estimated with the ZCA or GEV projections of the timeseries data, followed by classification analysis based on GC or PC connection strength as features. **C.** (Top) Two-way task versus resting state classification accuracies following ZCA-based decorrelation. Gray circles: Classification accuracies based on PC. Other conventions are as in Figure 1B. Dashed line: chance accuracy (50%).(Bottom) Same as in top panel, but for classification following GEV-based decorrelation. **D.** (Top) Schematic showing unweighted directed graph obtained from dGC; this digraph representation encodes only the dominant direction of connectivity, and not its magnitude. (Bottom) Two-way task versus resting state classification accuracies based on dGC digraph representations. Secondary ordinate (y-axis on the right): number of scan timepoints for each task.(Panels C-D). GC features were estimated with the Shirer et al [26] 14-network parcellation.

Second, we asked if lag-based connectivity could accurately classify task from resting state, once the data were purged of all instantaneous correlations. To accomplish this, we adopted two approaches: i) zero-phase component analysis (ZCA) and ii) generalized eigenvalue decomposition (GEV) (Methods). Briefly, ZCA (or the Mahalanobis transformation) produces whitened time series data that is closest, in a least squares sense, to the original regional time series data. As an alternative approach, we decorrelated both task and resting state time series jointly by projecting them onto a single set of generalized eigenvectors (GEV). These approaches provided empirical upper and lower bounds on GC’s performance on correlation-purged data (Methods).

GC connectivity features sufficed to successfully classify all tasks from resting state, even in correlation-purged data. With ZCA, iGC accuracies ranged from 84% to 96% whereas dGC accuracies ranged from 82% to 96% across tasks. With GEV, iGC accuracies ranged from 60% to 71% whereas dGC accuracies ranged from 56% to 76% across tasks; in each case, classification accuracies were significantly above chance (p<0.001, permutation test). We confirmed that performance in each case was not an artifact of the decorrelation procedure (ZCA/GEV) by randomly interchanging task and resting state labels for each pair of datasets across subjects (Methods); shuffling labels reduced classification accuracy to chance. Note that in every case, classification performance based on PC connectivity was at chance (Fig. 2C), a direct consequence of removing instantaneous correlations from the data. Despite this, classification accuracies based on iGC connectivity were not at chance; in the next section, we discuss potential reasons for these differences between iGC and PC classification accuracies.

Third, we asked if an unweighted directed graph (digraph) network representation – whose edges indicated the dominant direction, but not magnitude, of connectivity (Fig. 2D) – would suffice to distinguish task from resting brain states (Methods). Again, dGC directed graphs successfully distinguished each task from resting state well above chance. Classification accuracies ranged from 56% for the motor task versus resting state classification to 68% for the relational task versus resting state; for each task, classification accuracies were significantly above chance (p<0.001; permutation test). Interestingly, we did not see a strong influence of the number of data points on classification accuracy in this case (Fig. 2D, purple dots). For instance the emotion task (n=176 timepoints) was classified with an accuracy of 62% from resting state, which was comparable to the classification accuracy of working memory (n=405 timepoints) from resting state (64%). Both iGC and PC, which are symmetric connectivity measures, could provide no directed connectivity information.

These results demonstrate that lag-based connectivity contained sufficient information to classify task from resting state even when instantaneous correlations were entirely purged from the data. Moreover, unweighted directed connectivity graphs alone, indicating the direction, but not scalar magnitude, of GC connectivity, sufficed to accurately classify task from resting brain states. These findings indicate that directed functional connectivity measures, like dGC, provide connectivity information that is distinct from, and complementary to, what can be obtained with undirected functional connectivity measures, like PC.

### Instantaneous and directed GC identify complementary aspects of functional connectivity

What characteristics of functional connectivity are respectively identified by instantaneous and lag-based connectivity? And how can lag-based connectivity be reliably estimated with fMRI data, which is sampled at time scales orders of magnitude slower than neural timescales? We addressed both of these questions, first, with simulations (this section) and, then, with real data (next section).

First, we tested the ability of GC to reliably recover functional interactions in simple, two-node feedforward networks operating at different timescales (Fig. 3A). We simulated fMRI data using a two-stage model (Methods): i) a latent variable model that describes the dynamics of the nodes (vector Ornstein-Uhlenbeck process; [32]); ii) a convolution of these neural dynamics with a hemodynamic response function to obtain the simulated fMRI time series [18,19]. Based on this model, we simulated activity in two 2-node networks. In the first network, individual node decay timescales were set to 50 ms, whereas in the second network, these were set to 1000 ms (parameters in SI Table S6A). For convenience, we refer to these two network timescales as “fast” (50 ms) and “slow” (1000 ms). We then varied the sampling interval (T_s_) of the simulated data from 50 ms to 1450 ms in steps of 100 ms. Connections at both “fast” and “slow” timescales were generally discovered by iGC regardless of sampling interval, although connections at slow timescales were less robustly detected than those at fast timescales (Fig. 3A). On the other hand, the connection in the “fast” timescale network was not discovered by dGC when the sampling interval was higher than 50 ms, in line with the results of Smith et al [18]. However, the connection in the “slow” timescale network was reliably discovered by dGC across a wide range of sampling intervals, upto, and exceeding 1000 ms. In each case, dGC failed to discover the underlying interaction when the sampling interval was much higher than the slowest timescale in each network, consistent with recent theoretical results [6]. These findings suggest that dGC can detect slow neural processes, which operate at a timescale slower than TR, in fMRI data.

**Figure 3.**
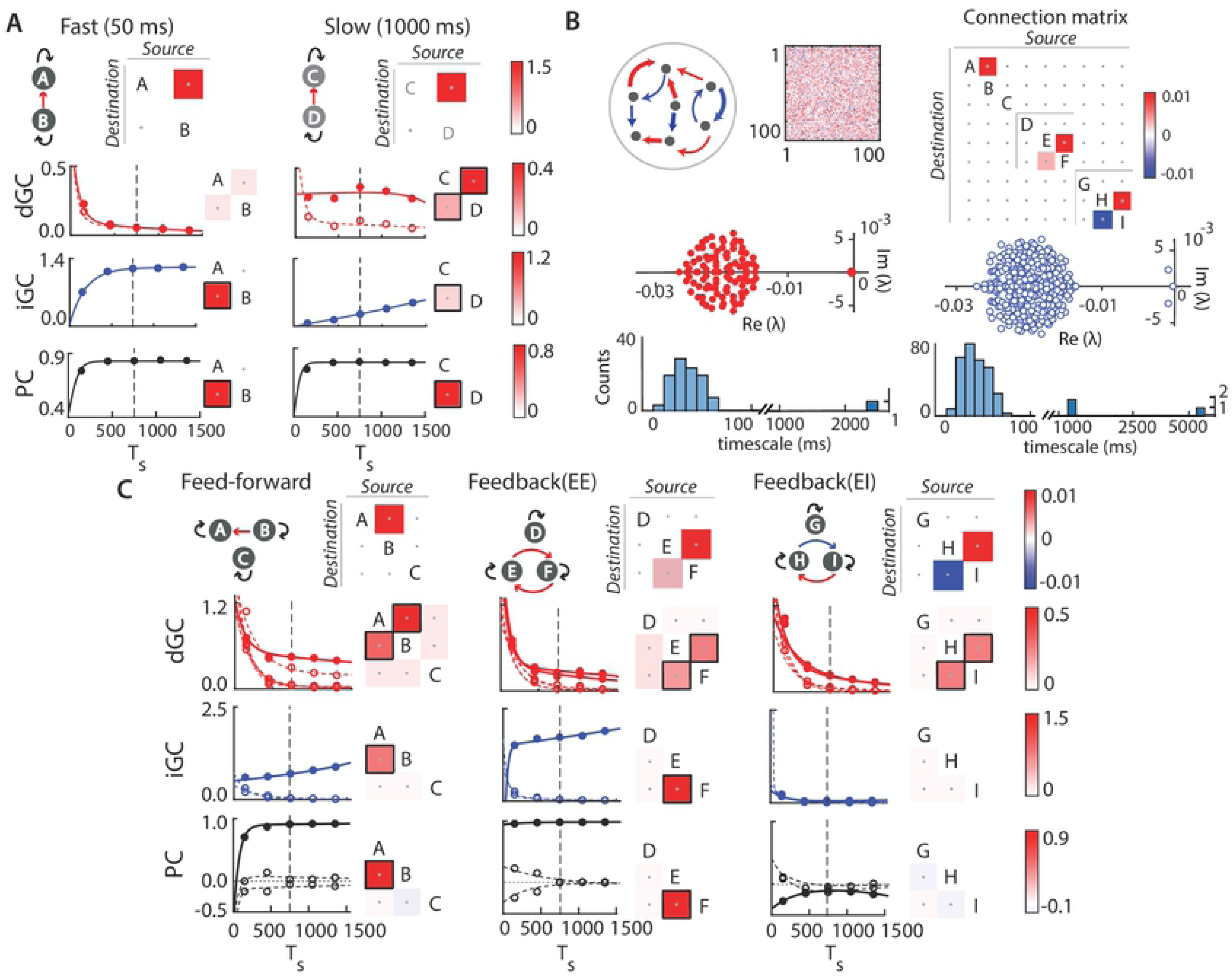
Robustness of GC estimates depend on network timescales in simulated hemodynamic data. **A.** (Top) Two-node networks with fast (50 ms; left) or slow (1000 ms; right) decay timescales of individual nodes. Each subpanel shows ground truth connectivity either as a schematic (left) or connectivity matrix (right). In the matrix, a non-zero entry at cell (i, j) corresponds to a connection from node j (source) to node i (destination).(Bottom) dGC (red), iGC (blue), and PC (black) connection strengths as a function of sampling intervals. Filled circles and solid lines: Strengths of true connections and curve (biexponential) fits, respectively. Open circles and dashed lines: Strengths of spurious connections and curve fits, respectively. Dashed vertical line: Sampling interval of 750 ms, mimicking the TR of the fMRI data. Matrices to the right of each plot show GC connection strengths estimated at sampling interval of 750 ms. Black squares surrounding matrix cells denote significant connections (Methods). For iGC and PC (symmetric connectivity), only the lower triangular matrix is shown, for clarity. **B.** (Top left) Schematic showing a cluster of neurons, each with timescale 50ms, connected with sparse, random, net excitatory connectivity. Matrix: Connectivity among the 100 neurons in a representative cluster. Red: excitatory connections; blue: inhibitory connections. Each such cluster forms one of the nine nodes in the simulated network. (Top right) Connectivity among the nine nodes in the network. (Bottom left) Eigenspectrum (upper panel) of a representative 100 neuron cluster, showing one slow emergent timescale corresponding to the real-part of one eigenvalue close to zero. Histogram (lower panel) showing timescales of all eigenmodes, with the slowest eigenmode at >2000ms. (Bottom right) Eigenspectrum (upper panel) of sub-network DEF exhibits multiple slow emergent timescales. Histogram (lower panel) showing timescales of all eigenmodes, with three slow eigenmodes at ∼1000-6000 ms. **C.** Same as in A, but for simulated 9-node networks. (Left) Sub-network ABC, (middle) sub-network DEF, (right) sub-network GHI. Other conventions are as in panel A.

How might such slow timescales, orders of magnitude slower than spike times and membrane time constants, arise in fMRI data? To answer this question, we availed of established results in random matrix theory. Connectivity in randomly connected E-I networks of neurons can produce slow timescales, without fine-tuning of network parameters [32,33]. We modeled sparse, random, net excitatory connectivity in a small network of (N=100) neurons with connection parameters drawn from previous studies (SI Table S6B; [32,34,35]). The eigenspectrum of the network revealed that each network exhibited one eigenvalue close to zero corresponding to a slow timescale (∼1000 ms or greater, Fig. 3B bottom left); the latter constitutes an emergent timescale associated with the dominant eigenmode that is a property of network connectivity (Methods).

We modeled nine such networks, organized into three, non-interacting, clusters (Fig. 3B top right): a) a cluster with a purely feedforward connection across two networks, b) a cluster with recurrent excitatory (E-E) feedback connections among two networks and c) a cluster with recurrent excitatory-inhibitory (E-I) feedback connections among two networks. In each case, connectivity across networks was mediated by a small proportion (5%) of neurons in each network (parameters in SI Table S6B). This configuration mimics “small-world” connectivity in brain networks [36], with locally-connected brain regions interacting through sparse, long-range connections [37]. The eigenspectra revealed that dynamics in all clusters operated at timescales of around 6000 ms, comparable to or slower than the individual network timescales (Fig. 3B bottom right). To simulate fMRI data we averaged the activity across all 100 neurons in each network and convolved it with a canonical HRF. As before, these nine timeseries were then sampled at various sampling intervals, including a 750 ms interval mimicking the scan TR, and analyzed with GC to detect significant connections.

iGC and dGC identified complementary aspects of connectivity with these simulated data (Fig. 3C). iGC robustly identified feedforward and excitatory (E-E) feedback connections. dGC also estimated these connections, albeit with the following differences: First, in the feedforward network dGC occasionally identified a spurious connection, albeit much weaker in magnitude, in the direction opposite to the true connection (Fig. 3C, left column, red dashed line). Second, when the E-E feedback connections were precisely balanced in strength (symmetric), dGC also failed to identify the connection reliably (SI Fig. S3A). Yet, when these connections were of different strengths dGC reliably identified both connections, and their relative strengths (Fig. 3C, middle column, red). In contrast, when the connections were of different signs (E-I feedback) dGC robustly identified both connections, whereas iGC failed to reliably detect this connection (Fig. 3C, right column, blue). Yet, taken together, iGC and dGC identified all three connection types reliably.

Next, we compared the efficacy of connectivity estimation with partial correlations (PC). While PC robustly identified both feedforward and feedback E-E connections (Fig. 3C left and middle columns, black), it, surprisingly, failed to estimate feedback E-I connections, particularly when these were balanced in strength (Fig. 3C right column, black). When the E and I connection strengths were not balanced, but were strongly biased in favor of the E or the I connection, PC estimates varied with the sign of the more dominant connection (SI Fig. S3B, right top). These results generalize beyond these particular simulations; in SI Mathematical Note, Section S2 and S3, we identify, analytically, network configurations for which PC estimates systematically deviate from ground-truth connectivity (see also SI Results, section on “Analytical relationship between PC and iGC”).

Taken together, these results indicate that instantaneous and lag-based connectivity measures can reveal complementary aspects of brain connectivity. In addition, the results challenge the notion that correlation-based measures, like PC, should be favored over lag-based measures, like dGC for measuring functional connectivity in the brain [18]. Rather, the strengths and weaknesses of each measure (GC and PC) must be recognized when seeking to apply these to brain imaging data.

### Identifying a cognitive core system and predicting behavioral scores with GC connectivity

Our classification analyses and simulations suggested that iGC and dGC reliably recover task-specific brain networks, the latter when slow-timescale processes occur within the network. We asked whether iGC and dGC connectivity merely reflected reliable statistical patterns of brain activity, or whether it would be relevant for understanding the nature of information flow in the brain, and its relationship to behavior. To answer this question, we first investigated whether each measure would identify brain networks with consistent outflow and inflow hubs across tasks. Next, we asked whether GC connectivity would be relevant for predicting brain-behavior relationship.

First, we sought to identify a common core of “task-generic” connections across cognitive tasks. For this, we applied a feature selection approach – recursive feature elimination (Methods) – a technique that identifies a minimal set of features that provide maximal cross validation accuracy (generalization performance) [38]. Prior to analysis of real data, we validated RFE by applying it to estimate connectivity differences in two simulated networks (Fig. 4A,B). RFE accurately identified connections that differed in simulation ground-truth: specifically, differences in fast timescale connections were reliably identified by iGC, and in slow timescale connections by dGC (Fig. 4B bottom). RFE based on dGC and iGC also accurately identified the relevant connections, but not always their directionality, even when systematic variations in hemodynamic lag occurred across regions; the results are described in SI Results, section on “Effects of regional variations in hemodynamic lag”.

**Figure 4.**
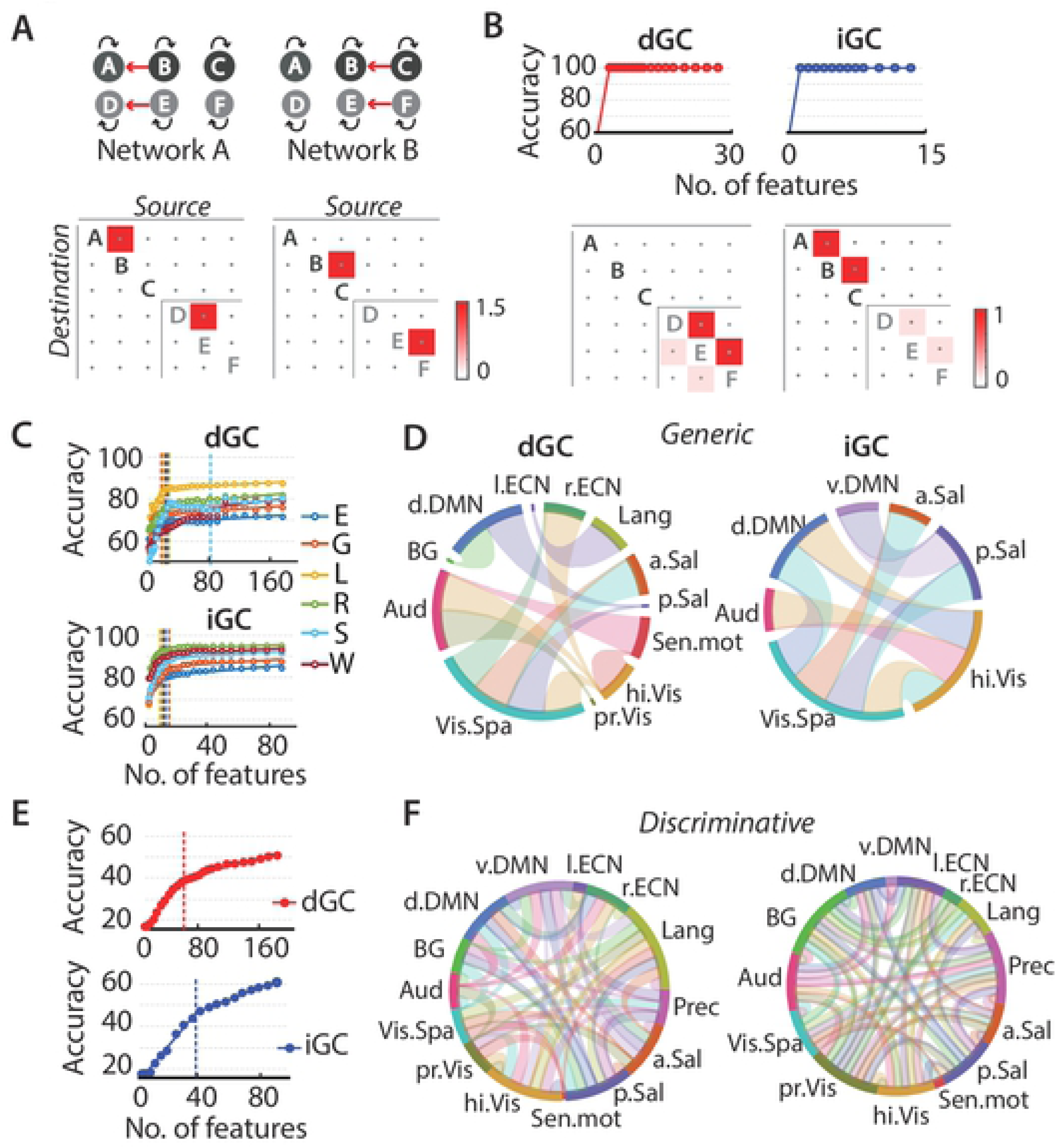
Recursive feature elimination (RFE) identifies task-generic and task-discriminative networks based on GC connectivity. **A.** Schematic showing two simulated networks each with fast (50 ms; ABC) and slow (1000 ms; DEF) sub-networks, with distinct connectivity patterns. Network activity was simulated for 375 seconds with a sampling interval of 5 ms, convolved with the hemodynamic response function and sub-sampled at 750 ms to yield 500 simulated time points. **B.** (Top) RFE curves, with classification accuracy as a function of remaining features, for classification based on dGC (left) and iGC (right). (Bottom) Maximally discriminative features following RFE based on dGC (left) and iGC (right). Entries denote average beta weights across RFE iterations. **C.** RFE curve for two-way classification of each of six tasks (all tasks except Motor) versus rest, based on dGC (top) and iGC (bottom). Color conventions are as in Figure 1D. Data points: RFE accuracies; solid lines: piecewise linear fits. Vertical dashed line: location of the elbow for each RFE curve. **D.** Task-generic connections following task-versus-resting RFE, based on dGC (left) and iGC (right) features, using Shirer et al (2012) 14-network parcellation [26] (SI Table S4); each network is indicated with a different color and a label. Directed dGC connections are shown as tapered links, broad at the source node and narrow at the destination node. Undirected iGC connections are shown as bidirectional links between the respective pair of nodes. Colors of the connections represent the color of the destination node. **E.** Same as in panel D, but for n-way classification across the six tasks. Color conventions are as in panel B. **F.** Same as in panel E, but for task-discriminative connections, which maximally discriminated each task from the five others, following n-way RFE, based on dGC features (left) and iGC features (right). Other conventions are the same as in panel C.

We applied RFE to classify tasks versus resting state; we chose these six tasks (all tasks except the motor task) as being the most likely to engage common cognitive control mechanisms (Fig. 4C). For these RFE analyses we employed a 14 network functional parcellation [26], as it consistently gave good classification accuracies with both iGC and dGC connectivity (SI Fig. S1B). Following RFE, we applied a binomial test across tasks (Methods) to identify a common core of task-generic connections, separately for iGC and dGC.

RFE identified distinct task-generic networks with iGC and dGC, which comprised of connections that distinguished a majority of tasks from resting state. The iGC task-generic network revealed a visuospatial network hub, which connected with the anterior salience, dorsal DMN, higher visual and posterior salience networks (Fig. 4D, right). The dGC task-generic network confirmed the hub-like connectivity of the visuospatial network but, in addition, revealed consistent directed information outflow from the visuospatial network to the other networks (Fig. 4D, left). In addition, dGC revealed consistent inflow into the higher-visual network across tasks, including from the visuospatial, right executive control, and auditory networks, consistent with the ability of top-down inputs from these networks to strongly modulate sensory encoding in higher visual cortex [39]. Finally, the higher-visual network projected consistently to the sensorimotor network, suggesting a final common pathway, across these tasks, for influencing behavior. Interestingly, the only network providing inflow into the visuospatial network hub was the anterior salience network, in line with a previous study that indicated a role for the salience network in controlling other task positive networks [7].

Similarly, we also identified connections that were maximally discriminative across tasks; again iGC and dGC showed distinct sets of these “task-discriminative” connections (Fig. 4E-F; SI Results, section on “Identifying task discriminative networks with GC”).

To address GC’s relevance for understanding brain-behavior relationships we tested whether the strength of functional connections estimated with iGC and dGC could predict inter-individual variations in behavioral scores as measured by a standard cognitive battery (Methods; SI Table S7). We employed a leave-one-out prediction analysis based on multilinear regression followed by robust correlations of predicted and observed scores (Fig. 5A; p<0.05 with Benjamini-Hochberg correction; Methods).

**Figure 5.**
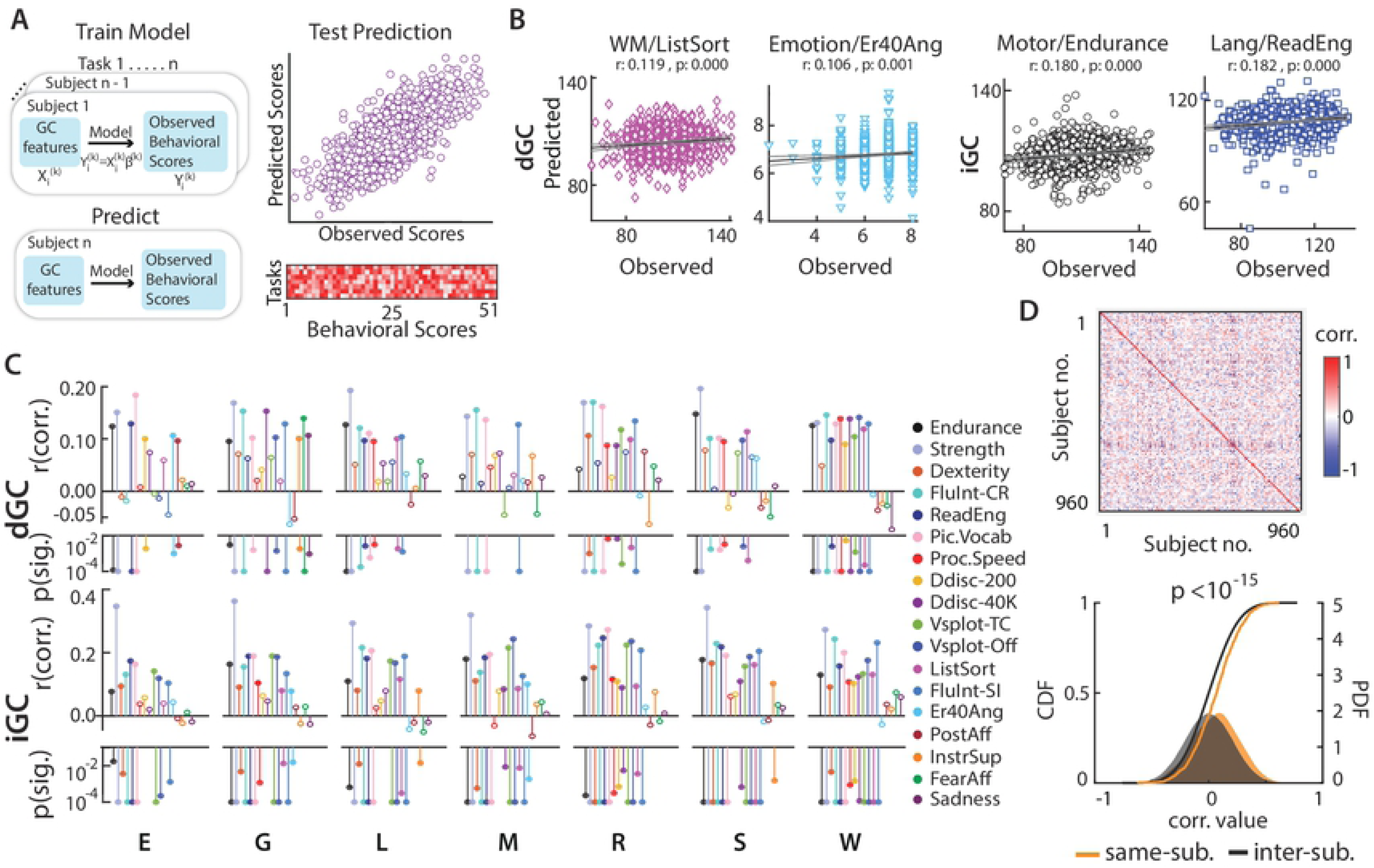
GC connectivity explains inter-individual variations in behavioral scores. **A.** (Left) Schematic of behavioral score prediction analysis. GC connectivity strengths for each task were used as independent factors to predict behavioral scores using linear regression with a leave-one-out approach. 51 different behavioral scores were predicted (SI Table S7), compared against observed scores (upper right), and their correlation values plotted as a matrix (lower right). **B.** Exemplar score predictions based on dGC (left panels) and iGC (right panels). In order (from left to right): List Sorting score predicted from Working memory task dGC connectivity, Anger Emotion Recognition score from Emotion task dGC connectivity, Endurance score from Motor task iGC connectivity and Reading score from Language task iGC connectivity. **C.** (Top) Prediction statistics for selected scores based on dGC connectivity. Correlation coefficients (r values) between the predicted and observed scores are plotted in the top half of each stem plot, and significance (p values) are plotted in the bottom half. Each score is denoted by a different color, and each sub-panel shows predictions based on GC connectivity for a different task; Stems with open symbols represent non-significant correlation coefficients, whose corresponding p-values are not shown. p values are floored at 10-4for ease of visualization. (Bottom) Same as in top panel, but predictions based on iGC connectivity. **D.** (Top) Inter-subject correlation matrix of composite behavioral scores. Row and column indices: subjects. (Bottom) Cumulative distributions (solid lines) and density function estimates (filled area) of correlation coefficients between observed and predicted composite scores, for the same subject (yellow) or across different subjects (grey). Predictions were based on GC estimates from the relational and working memory tasks. p-value: Kolmogorov-Smirnov test.

Both iGC and dGC predicted key behavioral scores. Several scores were predicted uniformly well by iGC across tasks (Fig. 5B, right; Fig. 5C, bottom; SI Fig S5B). Scores of fluid intelligence (Penn progressive matrices), grip strength, endurance, and language (reading and picture-vocabulary) (Fig. 5B right; r: 0.077 - 0.363; p<0.02), were all well predicted by iGC, in addition to scores of spatial orientation (Penn line orientation test)and dexterity (SI Fig. S5B, r:0.081 −0.243; p<0.0125). On the other hand, dGC-based predictions were more selective, in that several behavioral scores were best predicted by dGC based on specific tasks alone (Fig. 5B, left; Fig. 5C, top; SI Fig. S5A). For instance, dGC in the emotion task alone predicted positive affect (Fig. 5C, top; r=0.094, p=0.028) and anger emotion recognition (Fig. 5B, left, cyan; r=0.106, p=0.001) scores, dGC in the gambling task alone predicted self-report scores of perceived social support (r=0.101, p=0.002), fear (r=0.139, p<0.001) and sadness (r=0.107, p=0.001) and dGC in the motor task alone predicted median reaction time in the fluid intelligence test (r=0.123, p<0.001).In addition, dGC in the working memory task predicted a range of scores in the “cognition” category including list sorting(Fig. 5B, left, pink; r=0.119, p=0.000), fluid intelligence, picture discrimination speed, spatial orientation and self regulation (discounting of delayed reward; Fig. 5C top, SI Fig. S5A).

A variety of behavioral scores were also successfully predicted based on PC connectivity (SI Fig. S5C); several of these (∼60%) overlapped with those predicted with GC connectivity (SI Results, section on “Predicting behavioral scores with PC connectivity”). Yet, PC connection features that led to successful predictions overlapped strongly with iGC connection features than with dGC connection features, as quantified by the mean correlation between their regression weights in the respective prediction models (PC vs. iGC: r=0.38±0.01, mean±std; PC vs. dGC: r=0.03±0.01, p<0.001, ranksum test).

Finally, we tested whether GC connectivity could predict a combined set of behavioral scores unique to each subject. For this, we created a vector of all independent behavioral scores (composite score; Methods), and confirmed that this composite behavioral score uniquely identified each subject in the database, as evidenced by the highest values along the main diagonal of the inter-subject correlation matrix (Fig. 5D top). Following this, we performed the leave-one-out prediction, as before, except that we used dGC and iGC connectivity features from two of the tasks alone (working memory and relational; also see SI Fig. S5D). We then tested whether each subject’s predicted composite score would correlate best with her/his own observed composite scores. Although we did not observe the highest correlation values consistently along the main diagonal, the distribution of correlation coefficients along the diagonal were significantly different (and higher) than the distribution of off-diagonal correlation coefficients (Fig. 5D bottom; p<10^-15^, Kolmogorov-Smirnov test). Inter-individual variation GC connectivity, therefore, contained sufficient information to accurately identify subject-specific behavioral scores in this cohort of subjects.

In summary, the ability to successfully predict subject-specific behavioral scores suggests that GC functional connectivity is relevant for understanding brain-behavior relationships. Moreover, connection features that were relevant for behavioral predictions with PC overlapped highly with iGC, but not with dGC, thereby validating our simulation results regarding the complementarity of iGC and dGC connectivity estimates.

## Discussion

Neural processes in the brain range from the timescales of microseconds to milliseconds, for extremely rapid processes (e.g. sound localization), to timescales of several seconds to minutes, for processes that require coordination across diverse brain networks (e.g. when having a conversation), and hours to days, for processes that involve large-scale neuroplastic changes (e.g. when learning a new language). Coordinated activity among brain regions that mediate each of these cognitive processes should manifest in the form of functional connectivity among these regions at the corresponding timescales. Our results indicate that applying Granger-Geweke Causality (GC) with fMRI data permits estimating behaviorally relevant functional connectivity at a timescale corresponding to the sampling rate of fMRI data (seconds).

The application of GC to neuroscience is a contentious topic, for a variety of reasons [15–17,19,28]. One particular challenge stems from the use of the word “causality”: the notion of causality in GC is different from the notion of interventional causality [40]. Our use of the term Granger causality, here, purely reflects its application as a marker of information flow among brain networks [19,41], and is not meant to indicate causality in a physical, interventional sense.

With this understanding, our results contain three key insights. First, we show that, either iGC or dGC connectivity suffices to reliably classify task-specific cognitive states with superlative accuracies (Fig. 1B). Instantaneous and directed GC – both measures of conditional linear dependence and feedback [9] – were able to robustly estimate task-specific functional interactions even with slowly sampled fMRI data. Our application of machine learning and classification analysis circumvents the lack of access to ground truth connectivity, and our simulations suggest that GC connectivity is relevant for estimating slow, emergent interactions among brain networks [15–19].

Second, we show that functional connections identified by iGC and dGC carry complementary information, both in simulated and in real fMRI recordings, and we demonstrate key caveats with employing correlation-based measures of functional connectivity like partial correlations, despite superior classification accuracies with these latter measures. First, PC fails to correctly infer reciprocal excitatory-inhibitory interactions, which can be accurately inferred with lag-based methods like dGC. Second, PC may yield incorrect estimates of functional connectivity that do not match ground truth (SI Fig. S3C). In particular, when the data are well described by an autoregressive model framework our results suggest that instantaneous connectivity measures, like iGC, provide more accurate descriptions of functional connectivity than PC. Third, even with data completely purged of partial correlations, dGC connectivity was sufficient to classify task-specific cognitive states (Fig. 2C). In fact, unweighted directed connectivity alone sufficed to produce accurate classification at accuracies significantly above chance (Fig. 2D). These results indicate that information flow mapped by GC connectivity can be complementary to that of PC, and highlights the need for examining diverse measures, both instantaneous and lag-based, to obtain a complete picture of functional connectivity in the brain.

Third, differences in inter-individual iGC and dGC connectivity were able to successfully explain inter-individual variation in behavioral scores on various cognitive tasks, and to identify an individual-specific composite marker of behavioral scores, with high accuracies. Because these behavioral scores were acquired in a separate testing session outside the scanning session [27], the results suggest that GC connectivity was both individual-specific, and stable over timescales exceeding the scan session, to permit accurate prediction. Moreover, in our analysis, each subject’s behavioral score was predicted based on her/his GC connectivity, whereas the regression beta weights – representing the relationship between GC connectivity and behavior – were computed from the population of all subjects excluding that subject (Fig. 5A). Successful predictions, therefore, indicate a consistent mapping between GC connectivity and behavioral scores across the population of subjects. These findings complement recent results showing that dynamic, resting-state functional connectivity, based on correlations, can explain significant variance in human behavioral data [42], and indicate the relevance of lag-based connectivity measures for understanding brain-behavior relationships.

Does GC’s discriminatory power rely on directed functional connectivity in the underlying neural response or systematic distortions of this connectivity induced by subsampling [19] and hemodynamic filtering [20,21]? While our findings cannot completely rule out the latter hypothesis, we address, next, three key caveats raised by previous studies for estimating functional connectivity with fMRI-GC, and argue why our results support the former hypothesis.

First, several studies have shown that sub-sampling of neural time series, at the scale of fMRI TR, renders functional connections undetectable with GC [11,18–20]. In these studies, GC was estimated with simulated fMRI time series, sampled at an interval (TR) of seconds, and failed to recover underlying neural interactions, which occur at millisecond timescales (e.g. [18]). However, these claims depended strongly on the nature and timescale of the connectivity in the networks employed in these simulations. For instance, a widely cited study [18] employed purely feedforward connectivity matrices with a 50 ms neural timescale in their simulations, and argued that functional connections are not reliably inferred with GC applied to simulated fMRI data. In addition to being neurally implausible, such purely feedforward network configurations yield eigenmodes whose slowest timescales are identical with the timescales of individual nodes [43]. Therefore, such a configuration rendered lag-based measures like GC, irrelevant for estimating neural interactions from slowly sampled fMRI data [18,19]. Furthermore, such connectivity precludes the occurrence of slower, behaviorally relevant timescales of seconds, which readily emerge in the presence of feedback connections, both in simulations [32,33] and in the real brain [44,45]. Our simulations show that slow timescale interactions emerge in networks with sparse, random, net excitatory connectivity, mimicking connectivity in the neocortex [32,33,35].While earlier studies have employed large-scale, biologically plausible models [46,47] to demonstrate the emergence of slow (<0.1 Hz) emergent functional interactions among brain networks, our results build upon these previous findings and show that such emergent functional interactions at slow timescales can be readily inferred from simulated fMRI data with GC. In fact, GC connectivity continued to robustly classify distinct task states even when data were sampled at 2x or 3x the original sampling interval of the fMRI data. Thus, while it is likely that GC applied to fMRI data is unable to detect connections at timescales faster than TR, our results show that sufficient distinguishing information occurs in slow-timescale connections to enable accurate inter-task classification.

Sub-sampling alone, may also produce spurious GC causality; the precise conditions under which spurious GC arises for continuous time vector autoregressive processes is an area of active research, and must be addressed in future studies [6,20].

Second, previous studies have shown that systematic differences in hemodynamic (HRF) lags (e.g. time to onset, or time to peak) among brain regions may produce spurious dGC estimates [19,21]. With simulations we demonstrated that fMRI-GC could identify differences in slow-timescale network connectivity, despite systematic differences and heterogeneity in HRF onset latencies across nodes (SI Fig. S3D-E). In all cases, applying recursive feature elimination with either dGC or iGC features identified the precise subset of connections that distinguished distinct network configurations. In a majority of cases, dGC also correctly identified the directionality of these connections. In our simulations, the only scenario in which dGC features failed to identify the directionality of connections correctly, was when the onset latency in the “destination” nodes were biased to be systematically earlier than those in the “source” nodes. Nevertheless, in the real data it is unlikely that systematic inter-regional HRF differences were responsible for the observed superior classification accuracies. Variations in HRF delays would indeed confound dGC connectivity estimates – if they occurred consistently between brain regions across subjects and tasks (e.g. SI Fig. S3D, red curves). Yet, such a scenario cannot account for the high classification accuracies among tasks and sub-tasks based on dGC connectivity alone. In other words, even if HRF latency differences systematically biased dGC connectivity estimates, these estimates were sufficiently and reliably different across task cognitive states to permit accurate classification among them. Finally, network properties of key regions identified with fMRI-GC were consistent with their known functional properties of these regions. For instance, dGC identified the visuospatial network as an information outflow hub, across all six cognitive tasks (Fig. 4D left). The visuospatial network comprises frontal cortex regions, including the frontal eye field, as well as posterior parietal cortex, which are both widely implicated in visuospatial attention control [48–51]. In addition, the only network that provided task-generic incoming connections to the visuospatial network was the anterior salience network comprising the anterior fronto-insular cortex and the anterior cingulate cortex [52,53], regions implicated in feature-based attention and the suppression of distractors [54]. Information outflow from these key networks identified by dGC is consistent with their role in attention and executive control.

Third, simulations and theoretical results indicate that scanner noise can degrade, or even obliterate GC connectivity estimates [19]. On the other hand, our classification accuracies suggest that GC estimates were sufficiently robust to scanner noise to permit accurate task and sub-task classification in these data. In fact, we show that averaging dGC connectivity across as few as 5 subjects’ data improves classification accuracy to over 95%, for nearly all tasks (Fig. 1D). Such superlative classification accuracies are unlikely to have occurred if scanner noise were to significantly degrade GC estimates.

In sum, these results strongly indicate that slow functional interactions in the brain can be meaningfully inferred with GC from fMRI data. While the directionality of interactions measured by GC may need to be interpreted with care [11,21], our results suggest that fMRI-GC may be useful for formulating hypothesis about the role of particular brain regions in providing “top-down” control signals, for modulating activity in other brain regions [7,8], as well as for investigating the nature of information flow in cortical microcircuits with slow sampling rate techniques, such as calcium imaging [55]. The causal role of these brain regions in behavior can then be directly tested with interventional approaches such as transcranial magnetic stimulation, optogenetic inactivation or by examining patient populations with lesions in specific brain regions [56]. Such a systematic analysis will pave the way for a mechanistic understanding of how flexible functional interactions among brain regions mediate complex cognitive behaviors.

## Materials and Methods

### Ethics statement

The scanning protocol for the Human Connectome Project was approved by the Human Research Protection Office at Washington University at St. Louis’ (IRB # 201204036). Only de-identified, publicly released data were used in this study.

### Data and code availability statement

Data used in the study is available in the public domain at the Human Connectome Project database (https://db.humanconnectome.org/). Code used for analyses are available at the following link: https://figshare.com/s/9d9131a6780fc8197cf1

Data sharing permissions can be found at the HCP website. Code may be shared or re-used upon requesting the corresponding author. These data and code sharing policies comply with the requirements of all funding agencies supporting this research and comply with institutional ethics protocols.

### fMRI data, parcellation and time-series extraction

We analyzed minimally preprocessed brain scans of 1000 subjects, drawn from the Human Connectome Project (HCP) database (S1200 release) (age range: 22-35 years; 527 females); fMRI acquisition and preprocessing details are described elsewhere [23,25]. SI Table S2 shows the identifiers of the subjects from whom data were analyzed. Data were analyzed from resting state and seven other task conditions (SI Table S1): Emotion processing, Gambling, Language, Motor, Relational processing, Social cognition and Working memory; in most figures, these tasks are referred to with their initial letters. fMRI scans for the relational task were not available for 9/1000 subjects; therefore, we analyzed a total of 7991 scans across all tasks and subjects.

We employed five different brain parcellations based one anatomical atlas and four functional atlases (SI Table S3). For the tasks versus resting-state classification based on GC connectivity (first section of Results), all 5 parcellations were used. Based on the classification performance in this analysis, we picked the three parcellations with the highest accuracies (90 node and 14 network parcellations of [26] and 96 network parcellation of [57])and these were used for the pairwise classification analysis of each task versus the other as well as the n-way task classification analyses. Analysis with averaging GC features across subjects (Fig. 1D) was performed with a 90 node parcellation [26]. Classification analyses with data purged of instantaneous correlations and unweighted digraph representations (second section of Results) were performed with the Shirer et al [26] 14 network parcellations. Analyses involving identifying task-generic and task-discriminative networks, as well as behavioral score predictions, based on GC features (last section of the Results) were performed with the Shirer et al [26] 14 network parcellation. Voxel time series were extracted using Matlab and SPM 8 [58], and regional and network time series were computed by averaging the time series across all voxels in the respective region or network.

We employed parcellations with fewer, more coarse-grained regions, rather than fine-grained parcellations because Granger Causality estimates were more reliable when the number of regions was fewer than the number of timepoints. Both task and resting scans were of sufficient duration (∼200-300 volumes) to permit robust GC estimation. Finally, we noticed that in some parcellations, there were overlapping voxels between some of the regions. To avoid mixing of signals, we assigned each overlapping voxel to the region to whose centroid it was closest, based on Euclidean distance.

### Estimating functional connectivity with GC

We modeled instantaneous and lag-based functional connectivity between brain regions using conditional Granger-Geweke Causality [9]. The linear relationship between two multivariate signals **x** and **y** conditioned on a third multivariate signal **z** can be measured as the sum of linear feedback from **x** to **y** (Fx→y|z), linear feedback from **y** to **x** (Fy→x|z), and instantaneous linear feedback (Fx◦y|z) [9,41]. To quantify these linear relationships, we model the future of each time series in terms of their past values, using multivariate autoregressive (MVAR) modeling (SI Mathematical Note, Section S1, equation 1). MVAR model order was determined with the Akaike Information Criterion (AIC) for each subject, and was typically 1. The MVAR model fit was used to estimate both an instantaneous connectivity matrix using iGC (Fx◦y|z) and a lag-based connectivity matrix using dGC (Fx→y|z). Details are provided in SI Mathematical Note, Section S1.

Briefly, Fx→y|z is a measure of the improvement in the ability to predict the future values of **y** given the past values of **x**, over and above what can be predicted from the past values of **z** and **y**, itself (and vice versa for Fy→x|z). Fx◦y|z, on the other hand, measures the instantaneous influence between **x** and **y** conditioned on **z** (see SI Mathematical Note, Section S1). We refer to Fx◦y|z, as *instantaneous* GC (iGC), and Fx→y|z and Fy→x|z as lag-based GC or *directed* GC (dGC), with the direction of the influence (**x** to **y** or vice versa) being indicated by the arrow. The “full” measure of linear dependence and feedback Fx,y|z is given by: Fx,y|z = Fx→y|z + Fy→x|z + Fx◦y|z. Fx,y|z measures the complete conditional linear dependence between two time series. If, at a given instant, no aspect of one time series can be explained by a linear model containing all the values (past and present) of the other, Fx,y|z will evaluate to zero [41].

### Classification with linear SVM based on GC connectivity

The connection strengths of the estimated GC functional connectivity matrices were used as feature vectors with a linear classifier based on SVM for high dimensional predictor data. For a parcellation with n ROIs, the number of features for iGC-based classification was n(n-1)/2 (upper triangular portion of the symmetric n×n iGC matrix) and for dGC-based classification it was n^2^−n (all entries of the n×n dGC matrix, excluding self-connections on the main diagonal).Based on these functional connectivity features, we asked if we could reliably distinguish each task condition from resting state (e.g. language versus resting) or each task condition from the other

For pairwise classification of resting state scans versus each task we used Matlab’s `fitclinear` function, optimizing hyperparameters using a 5-fold approach: by estimating hyperparameters with five sets of 200 subjects in turn, and measuring classification accuracies with the remaining 800 subjects. Classification performance was assessed with leave-one-out and 10-fold cross-validation. We also assessed the significance of the classification accuracy with permutation testing (see Methods). In simulations, we observed that the magnitude of GC estimates varied based on the number of timepoints used in the estimation. To prevent this difference in number of timepoints from biasing classification performance, each scan was truncated to a common minimum number of time samples across the respective scans being classified (task, resting) before estimating GC. For each subject, GC connectivity was estimated independently for the two scan runs (left-to-right and right-to-left phase encoding runs), and averaged across the runs. Hyperparameters optimized included the regularization parameter, regularization method (ridge/lasso) and the learner (linear regression model, svm/logistic) using the `OptimizeHyperparameters` option to the `fitclinear` function. Hyperparameter optimization was performed only for task vs. rest classifications, but not for subject feature averaging, task vs. task, or N-way classification analyses.

For pairwise classification of each task versus the other, default hyperparameters were used in the `fitclinear` function and classification performance was assessed with leave-one-out cross-validation. For n-way classification, we used MATLAB’s `fitcecoc` function, which is based on error-correcting output codes, and fits multiclass models for SVMs. Briefly, the function implemented a one-vs-all coding design, for which seven (number of classes in multiclass classification) binary learners were trained. For each binary learner, one class was assigned a positive label and the rest were assigned negative labels. This design exhausts all combinations of positive class assignments. Classification performance in n-way classification was assessed with leave-one-out cross-validation. For each classification analysis mentioned above, task scans were truncated to the common minimum number of time samples across each set of scans, before estimating GC.

### Classification based on GC connectivity across sub-tasks and with sub-sampled data

Tasks in the HCP data were run as a block design, alternating between various conditions (sub-tasks). We tested whether GC connectivity would be able to classify among sub-tasks within each task (SI Table S1B). fMRI time series corresponding to each sub task was obtained by concatenating blocks of fMRI task time series pertaining to the respective sub task; the temporal order across blocks was preserved while concatenating the data. We also ensured that data at the conjunction of two successive blocks, which represented non-contiguous time points, were not used for GC estimation. The two sub tasks to be classified were then truncated to have same number of time points. GC estimation and pair-wise classification across sub-tasks was performed with the procedure described in the previous section. The Shirer et al [26] 14-network parcellation was used for these analyses. For the motor task, time series for the left and right finger movement blocks were combined into a “hand” movement sub-task, and left and right toe movement blocks were combined into a “foot” movement sub-task.

We also tested whether GC on fMRI data sampled at slower rates would suffice to classify among task and resting states. We obtained time series downsampled at 2x the original sampling interval by removing data at even numbered sample points, and retaining data at odd numbered sample points (k=1, 3, 5…). The even-sample point data were appended the end of odd-sample data series, thereby retaining the overall number of data points in the original time series. Again, we ensured that data at the conjunction of the odd- and even-sampled data series (last odd-sampled point and first even sampled point), which represented non-contiguous data points, were not used for GC estimation. Similarly, we obtained time series downsampled at 3x the original sampling interval by removing every third data point, starting with the second or third data point, and concatenating these timeseries to retain the overall number of data points in the original timeseries. As before, GC estimation and pair-wise classification was performed with the procedure described in the previous section

### Permutation testing of classifier accuracies

We performed permutation tests for evaluating the statistical significance of classifier performance, using the method outlined in [59]. The test involved permuting task labels independently for each subject and computing a null distribution of 10-fold cross-validation accuracy. We employed 1000 surrogates and assessed significance of each empirically estimated 10-fold cross-validation accuracy values for dGC and iGC, based on the proportion of samples in the null distribution which were greater than the cross-validation accuracy estimated from the data. We conducted these analyses for the tasks versus resting state classifications, n-way task classification, classification analyses after purging instantaneous correlations and those based on digraph features, separately for the two metrics (dGC and iGC).

### Testing for data stationarity and goodness of MVAR model fit

Computing GC based on VAR modeling assumes that the timeseries represent a stationary process. Four different tests were performed to test whether the MVAR model provided a valid and adequate fit to the data (SI Table S5). We performed these tests for parcellated time-series using scripts provided in the Multivariate Granger Causality (MVGC) toolbox [60]. First, we checked for the stability of the MVAR model fit by computing logarithm of the spectral radius using the *var_specrad()* function. A negative value was taken to indicate a stable fit. Second, we assessed consistency of the model fit, which quantifies what proportion of the correlation structure in data is accounted for by the VAR model, using the *consistency()* function. We adopted a threshold of 80% (or above) for both task and resting timeseries to consider the data to have passed the test for consistency [60]. Third, we evaluated the whiteness of residuals based on the Durbin-Watson test for absence of serial correlation of VAR residuals, using the *whiteness()* function. Values of the Durbin-Watson statistic less than 1 or greater than 3 signify a strong positive or negative correlation, respectively among the residuals [60]. Subjects for whom the Durbin-Watson statistic lay between 1 and 3 for more than 90% of the regional timeseries, for both task and resting state data, were considered to have passed the test. Fourth, we checked for stationarity based on the augmented Dicky-Fuller unit-root test (ADF), using the *mvgc_adf()* function. As in the previous case, subjects for whom the ADF test statistic was less than its critical value for more than 90% of the regional timeseries, for both task and resting state data, were considered to have passed the test.

### Control for motion artifacts

We checked whether systematic differences in motion artifacts could contribute to the superlative classification accuracies observed with GC. For this, we calculated Frame-wise Displacement (FD) [30] as the sum of temporal derivatives of translational and rotational displacement along the three (x,y,z) axes in mm, with the estimated motion parameters provided by HCP. Frames with FD>0.5mm were considered “misaligned” and were discarded (“scrubbed”) while estimating GC values. Because dGC is estimated based on lagged correlations, we also discarded one frame before and after every misaligned frame (AR model order was typically 1 for these data). We then repeated the SVM-based two-way classification of resting state from the seven different task states, with GC features estimated on the “motion scrubbed” data; we also repeated n-way classification among the 7 tasks. Comparison of classification (cross-validated) accuracies with and without motion scrubbing, across all 1000 subjects, is shown in SI Fig. S2C.

### Functional connectivity estimation and classification with partial correlations

We compared the performance of classification based on GC measures with that based on partial correlations (PC). Partial correlations were computed based on the inverse of the covariance matrix as outlined previously [4,5]. Like iGC, the PC connectivity matrix is undirected and symmetric. Therefore, only the upper triangular portion of the matrix, including (n*(n−1)/2) PC weights, was used as features in the classification analyses. Classification and cross-validation analyses followed the procedures described in the Methods section on “*Classification with linear support vector machines based on GC connectivity”*.

PC connectivity performed consistently better than GC connectivity for classifying task from resting state (Fig. 2A). We propose the following analytical explanation for this observation: PC, an estimator based on instantaneous covariance, is less susceptible to noise than GC, which is based on lagged covariance. This is due to the fact that the estimation of lagged-covariance is susceptible to errors from noise at multiple time-points. For illustration, consider a timeseries generated by a VAR(1) model: **x**(t) = *A* **x**(t − 1) + **e**(t). The lagged (lag-1) covariance matrix (Σ_1_) is estimated from the data as:

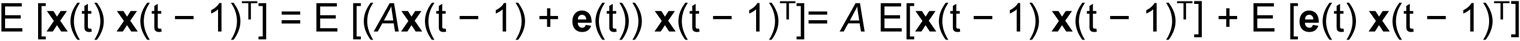

Thus, when estimating the lagged covariance, the variance of the interaction term E [**e**(t) **x**(t − 1)^T^] (second term in the right hand side) contributes to the variance of Σ_1_ in addition to the variance in computing the instantaneous covariance E[**x**(t − 1) **x**(t − 1)^T^] (first term on the right hand side).

### Classification based on GC connectivity in zero-lag correlation purged data

To test the complementarity of information conveyed by GC functional connectivity versus functional connectivity based on instantaneous correlations we decorrelated the regional time series data to purge them of instantaneous correlations. We adopted two approaches for this purpose: i) zero-phase component analysis (ZCA) and ii) generalized eigenvalue decomposition (GEV).

i. *Zero-phase component analysis (ZCA).* Consider demeaned *t*×*r* data matrix **X** of regional timeseries with *t* timepoints and *r* regions, with covariance matrix **C**. Decorrelating the data, to remove correlations among the columns of **X**, is achieved with a whitening transformation. A common whitening transformation is based on principal components analysis (PCA): **Y = W_PCA_X**, with **W_PCA_ = D^−1/2^E^⊤^**where **D** is a diagonal matrix, with the eigenvalues of **C** on its diagonals, and the columns of **E** contain the eigenvectors of **C**. While the PCA transformation effectively decorrelates regional timeseries, there is no way to ensure one-to-one correspondence of the whitened dimensions across subjects, rendering subsequent classification analysis challenging. Consequently, here we chose a different whitening transformation based on zero-phase component analysis (ZCA), also known as the Mahalanobis transformation. Based on this transformation, whitening is achieved as: **Y = W_ZCA_X,** with **W_ZCA_ = ED^−1/2^E^⊤^ = C^−1/2^**. A particular advantage of the ZCA transformation is that it yields whitened data that is as close as possible to the original data, in a least-squares sense [61]. Therefore, each subject’s data is projected on to a set of dimensions are most closely aligned with the underlying regional timeseries dimensions. Because the regions exhibit spatial correspondence across subjects (due to fMRI spatial normalization), the ZCA dimensions possess a natural, one-to-one correspondence across subjects, permitting subsequent classification. Before classification analysis ZCA dimensions were identified for each subject, separately for task and resting datasets. Regional time series for task and resting data were independently decorrelated by projecting onto their respective ZCA dimensions. GC (and PC) functional connectivity was estimated based on the these decorrelated timeseries, followed by classification analysis, as described previously (Methods section on “*Classification with linear support vector machines based on GC connectivity”*). As proof that the ZCA transformation was working effectively, classification accuracy based on PC (an instantaneous correlation measure) computed from ZCA components was at chance across all tasks (Fig. 2C top).
ii. *Generalized Eigenvalue Decomposition (GEV).* Although ZCA effectively purged correlations from the data, for the subsequent classification analyses task and resting state data were projected onto different, respective ZCA dimensions. Thus, the above-chance task versus resting state classification accuracy with GC features derived from ZCA components (Fig. 2C top) could perhaps be explained by, for example, systematic differences with how reliably ZCA dimensions were estimated across task and resting-state scans. We therefore sought an approach that could project both task and resting data into the same dimension while simultaneously decorrelating both. Such joint decorrelation may be achieved by projecting the data on to the generalized eigenvectors of the covariance matrices of the two datasets [62]. Let **C_T_** and **C_R_** denote the covariance matrices of the task and resting datasets respectively. The generalized eigenvectors of these two symmetric matrices are given by the columns of **G** = **E_T_ D_T_^-1/2^ E_R_,** where, as before **D_T_** is a diagonal matrix, with the eigenvalues of **C_T_** on its diagonals, and the columns of **E_R_** and **E_T_** contain the eigenvectors of **C_R_** and **C_T_** respectively. It can be readily verified that **G**^T^**C_T_G** and **G**^T^ **C_R_ G** are both diagonal matrices. Therefore, **G** is a matrix that jointly diagonalizes both **C_T_** and **C_R_**and projecting either task or resting state data into the columns of G decorrelates the respective timeseries. So, for these analyses, the regional time series for the task and resting state conditions for each subject were jointly decorrelated by projecting them onto a single space spanned the generalized eigenvectors. This was followed by classification analysis with GC features obtained from the decorrelated time series. As before, we confirmed the effectiveness of the decorrelation by computing classification accuracy based on PC from GEV components, which was at chance across all tasks (Fig. 2C bottom).

### Classification based on unweighted digraph representations of GC connectivity

An unweighted directed graph (digraph) network representation shows the dominant direction (but not magnitude) of functional connectivity among brain regions. Obtaining significant directed connections with dGC is challenging due to number of multiple comparisons required for testing n^2^-n connections. To identify significant directed connections, overcoming the multiple comparisons problem, we first subtracted the dGC connectivity matrix from its transpose and then applied the following two-stage procedure. In the first stage, the 1000 subjects were divided into five folds. For each two-way task versus resting state classification, recursive feature elimination (RFE, described in a later section titled “*GC feature selection based on Recursive Feature Elimination***”**) was performed based on dGC features of subjects from one fold (i.e. with 200 subjects). A minimal set of connection features identified by RFE, and their corresponding symmetric counterparts were then employed in the subsequent analyses; we term these connections K; the cardinality of K (the number of significant connections) was typically in the range of 2 - 86 (2.5^th^ - 97.5^th^ percentile). In the second stage, we identified statistically significant connections among these K features alone. For each of the subjects in the four remaining folds (i.e. 800 subjects), a null distribution for the dGC values of the features in K was obtained by estimating dGC following phase-scrambling the time series [8]. Next, we identified significant connections based on dGC values that occurred at the tail of the null distribution; the threshold for significant connections was determined based on a p-value of 0.05 with a Bonferroni correction for multiple comparisons. Classification performance based on digraph features was assessed with leave-one-out cross-validation.

### GC connectivity in simulated fMRI time series

To test the ability of GC measures to reliably recover functional interactions at different timescales, we simulated fMRI time series for model networks. Simulated fMRI time series were generated using a two-stage model. The first stage involved modeling latent neural dynamics with a stochastic, linear vector differential equation given by:

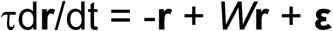

where **r** is the multivariate neural state variable representing the state of each neuron (or node) in the network (an N×1 vector, with N being the number of neurons), d**r**/dt is its temporal derivative, *W* is the neural (“ground truth”) connectivity matrix (dimension N×N), τ is the time constant of each neuron (or node) and **ε** is i.i.d Gaussian noise (N(0, Σ)), with Σ=*I*_N_ (N×N identity matrix). Although this model does not explicitly incorporate signal propagation delays, such vector Ornstein-Uhlenbeck models rank, arguably, among the most common models employed for simulating neural and fMRI time series, in many previous studies [6,18,19].The multivariate time series **r**(t), sampled at discrete time points r(kΔ) with a sampling rate of Δ, were generated based on the discrete time (1-lag) connectivity matrix A(Δ) and a residual noise intensity Σ(Δ). Here:

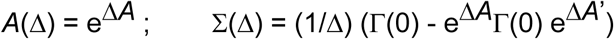

where *A* = (1/τ) (*W* - *I*_N_), e*^A^* denotes the matrix exponential, *A*’ is the transpose of *A*, and Γ(0) is the zero lag autocovariance which satisfies the continuous time Lyapunov equation *A*Γ(0)+Γ(0)A’+Σ=0 [19]. In the second stage, the latent neural dynamics were convolved with the hemodynamic response function (HRF) to obtain the simulated fMRI time series: **y** = *H* ⊗**x,** where *H* is the canonical hemodynamic response function (hrf; simulated with *spm_hrf* in SPM8), ⊗ is the convolution operation and **y** is the simulated fMRI time series. Finally, following convolution with the hrf, the data were downsampled to 750 ms, to mimic the repeat time (TR) of the HCP fMRI scans used in this study. The same model was used for the different simulations used in the manuscript (third section of the Results). The parameters for the 2-node simulations, and for the 9-node (100 neurons per node) simulations are described in SI Table S6.

For the 2-node simulations, iGC and dGC values were estimated by simulating the network for 200 timepoints, averaged across 25 repetitions. The 9-node simulations were performed with a 900 neuron network, with 100 neurons per node. Each node had sparse, random excitatory/inhibitory connectivity among its neurons (parameters in Table S6), whereas only 5% of neurons in each node were involved in inter-node connections, to mimic sparse, long-range connectivity in the neocortex [63]. The network was simulated for 200 timepoints, and timeseries from all (100) neurons in each node were averaged to generate 9 node timeseries. iGC and dGC values were estimated from the node timeseries and averaged across 10 independent repetitions. Significance was assessed with a bootstrap approach that involved generating 1000 surrogates by phase scrambling the node timeseries to yield a null distribution of GC values [8], followed by a Benjamini-Hochberg correction for multiple comparisons.

Simulations comparing PC and iGC connectivity were performed as follows: We simulated a 7-node network with a 1-lag VAR model of the form: **X**_k_ = *A* **X**_k−1_ + **ɛ**_k._ where **X**_k_ is the state of the discrete time process at discrete timestep ‘k’, *A* is the connectivity matrix, and **ɛ** is Gaussian noise with covariance matrix Σ_d_. *A* was chosen to be a random matrix with spectral radius less than 1 to ensure stability. Σ was chosen such that the covariance between every pair of residuals was zero (independent residuals) except for the first two residuals. The correlation between these residuals, ɛ^1^ and ɛ^2^, was parametrically varied between - 1.0 and 1.0 to systematically vary the strength of iGC connectivity. Note that, under this model, iGC between X^1^ and X^2^ vanishes only if and only if ɛ^1^ and ɛ^2^ are uncorrelated [9].

### GC feature selection based on Recursive Feature Elimination (RFE)

We performed features selection for analyses reported in Fig. 2D, 4B,C and S4B based on Recursive Feature Elimination (RFE). RFE identifies a minimal set of features, which provide maximal cross-validation accuracy [38]. Here, we implemented a two-level algorithm, described previously [16,64]. First, the data were divided into N_1_ (here, 10) folds. Of these, N_1_−1 folds were used as “training” data, and one fold was reserved as “test” data for quantifying the generalization performance of the classifier. Training data were pooled and further divided into N_2_ (here, 5) folds. The SVM classifier was then trained on N_2_−1 folds (leaving out one fold) and discriminative weights were obtained. The above procedure was repeated N_2_ times by leaving out each fold, in turn. Average weights were then computed by averaging the absolute values of the discriminative weights across the N_2_ runs. Next, 10% of the features (connections) contributing the lowest average weights were discarded, and the classifier was trained again with only the retained set of features. This procedure of feature selection and training was repeated until no more features remained. At this stage, the generalization performance for every set of retained features (each “RFE level”) was assessed using the left out “test” data. The entire procedure was repeated N_1_ times by leaving out each fold of the original data, in turn, as test data. Final generalization performances and discriminative weights of each RFE level were obtained as the average over N_1_ folds. We selected the set of connections at the RFE level at which the generalization performance reached an “elbow”: a minimal set of connections at which generalization performance dipped dramatically below its maximal level. To identify this elbow (e), we used a custom elbow fitting procedure, requiring a piecewise linear fit to the RFE curve, based on two lines, one for “x>e” and another for “x<=e”, with the first line required to have a higher slope than the second. The first point in each RFE curve was excluded from the higher slope line fit (Fig. 4C, 4E, SI Fig. S4B). RFE was typically repeated 5 times before determining peak accuracy and corresponding features.

### Simulating hemodynamic lag variations across nodes

We simulated systematic differences in hemodynamic lags across nodes by varying the onset parameter of the *spm_hrf* function (SPM8; [58]). For network configurations A and B described in Figure 4A, we simulated 4 scenarios: a) same mean HRF onset (μ_L_= 3s) across nodes; b) source node HRF onset lagging the destination node by 1s (μ_L-src_ > μ_L-dst_); c) source node HRF onset leading destination node by 1s (μ_L-src_ > μ_L-dst_); and d) mixed latencies of lead and lag across source and destination nodes (see next). GC was estimated for 100 simulated participants, by sampling onset latencies for each of the 6 nodes (A-F) from normal distributions (truncated to have only positive latency values), over a range of different standard deviations (σ_L_=0-1s, in steps of 0.2s). Onset latencies were sampled independently across participants, but were sampled such that the relative latency between each pair of source and destination nodes, across corresponding network configurations, remained the same for each participant. For example, if the onset latency difference between nodes A and B was 0.7s (μ_L-B_−μ_L-A_=0.7s) for a particular subject, the same difference in onset latency was also maintained between nodes B and C (μ_L-C_−μ_L-B_=0.7s). For simulations with mixed latencies (case d), 50% of simulated participants had onset latencies drawn from distributions with the source node lagging the destination node (case b) and the remaining 50% with the source node leading the destination node (case c). GC values were averaged over 5 runs for each simulated participant. Finally, we performed RFE to identify key connections that distinguished the two network configurations (same procedure as in Fig. 4B). Connections weights of the most discriminative connections following RFE are shown in SI Fig. S3E (for σ_L_=0.4s). Difference of dGC connections strengths as well as iGC connection strengths, for various values of σ_L_, are shown in SI Fig. S3D.

### Identifying “task-generic” and “task-discriminative” GC connections

To identify a minimal set of connections that occurred consistently across tasks (“task-generic” connections), we adopted the following approach. We performed RFE analysis for task versus resting state classification for each of the six tasks (all tasks except motor); we expected each of these tasks to recruit common cognitive control mechanisms. We then performed a binomial test to identify connections that were consistently activated across tasks. Briefly, the presence or absence of a connection in the set of RFE features for a given task versus resting state classification was considered as a Bernoulli trial, with probability of success (its presence) *p* being the mean number of RFE features identified across all six classifications. The number of trials *n* was the number of tasks versus resting state classifications (here n=6). The probability of a randomly picked connection being present in more than *k* such RFE sets is given by the cumulative distribution function for the binomial distribution *F(k; n, p).* Significant connections were identified as those that occurred in *k* or more tasks, with threshold at the p=0.05 level.

To identify a minimal set of connections that maximally differed across tasks (“task-discriminative” connections), we used RFE with an n-way classifier, to classify among all six tasks (again, except the motor task). The n-way classifier is based on training *n* (here, 6) one-vs-all binary learners. At the second level of the RFE procedure described above, average weights were computed for each of these *n* binary learners by averaging the absolute values of the discriminative weights across the N_2_ runs. Next, a set of features obtained by taking union of 1% of the features (connections) contributing the lowest average weights in each learner was discarded, and the classifier was trained again with only the retained set of features.

While quantifying the overlap between task-generic and task-discriminating connections identified separately for dGC, iGC and PC, we converted the dGC matrix to a lower triangular matrix by reflecting all connections about the main diagonal. The degree of overlap between PC and GC connections was quantified as the number of overlapping connections as proportion of the total number of connections identified by PC. We then computed a null distribution of the degree of overlap by randomly permuting the connection identities within each matrix, while preserving the overall number of connections in each matrix, and generating 1000 surrogate samples. The significance of the overlap of task-generic or task-discriminating connections between each pair of metrics (PC-dGC or PC-iGC) was quantified as the fraction of overlapping connections in the data that exceeded this null distribution.

### Predicting behavioral scores based on GC connectivity

We asked whether inter-individual differences in GC connectivity would be relevant for predicting inter-individual differences in behavioral scores. HCP provides a well-validated battery of behavioral scores assessed with a wide range of cognitive tasks. The task battery is based on the NIH Toolbox for Assessment of Neurological and Behavioral function [65], developed to create a uniform set of measures for rapid data collection in large cohorts. The toolbox includes assessments of cognitive, emotional, motor and sensory processing scores in healthy individuals. We pre-selected, based on domain knowledge, a specific subset of 51 scores for these analyses, using age-adjusted scores, wherever available (listed in SI Table S7). Next, we sought to predict subjects’ behavioral scores based on GC connectivity with an established leave-one-out approach [66]. Briefly, we used linear regression to predict behavioral scores using, as features, GC estimates of functional connectivity, separately for iGC (91 features or connections) and dGC (182 features). The leave-one-out analysis was performed such that the support vector regressor was fit on all but one subject and the learned beta weights were used to obtain predictions of the left-out subject’s behavioral score, using that subject’s own GC connectivity weights. Predicted scores were correlated with the actual scores using robust correlations (“percentage-bend” correlations; [67]).

Next, we asked if GC connectivity could identify an individual based on a composite marker of her/his behavioral scores. Because 40 subjects did not have a full complement of behavioral scores, data from the remaining 960 subjects was included in this analysis. The 51 behavioral scores were, each, z-scored across subjects and formatted into a “composite behavioral score” vector. This vector served as an individual specific composite marker of behavioral scores, as revealed the weak off-diagonal values in the covariance matrix of this vector across subjects (Fig. 5D top). dGC and iGC features of individual tasks, as well as combination of tasks (Relational and Working memory), were used to then predict the composite score marker for individual subjects, using the same leave-one-out procedure as described above. The observed and predicted set of composite scores was correlated across subjects. The distribution of observed versus predicted correlation values for each subject (values on main diagonal; Fig. 5D bottom yellow) were compared against between-subject correlation values (off-diagonal values; Fig. 5D bottom grey) using a Kolmogorov-Smirnov test.

## Acknowledgments

The authors would like to thank Lionel Barrett and Catie Chang for their comments on a preliminary version of this manuscript, and Govindan Rangarajan and Arshed Nabeel for helpful discussions.

## Supporting Information

**SI Text. Results and SI Figure Legends**

**SI Figure S1. GC classification accuracies for different parcellations and alternative classifiers**

**SI Figure S2. Stationarity tests and 1-stage versus 2-stage GC**

**SI Figure S3. Relationship between network connectivity, GC and partial correlations**

**SI Figure S4. Task generic and discriminative connections based on partial correlations (PC)**

**SI Figure S5. Behavioral score predictions based on GC connectivity strengths**

**SI Table S1A. Task descriptions.**

**SI Table S1B. Description of sub-tasks**

**SI Table S2. Subject identifiers**

**SI Table S3. Parcellations used in the analysis**

**SI Table S4. Network labels in the Shirer et al (2012) 14-network parcellation**

**SI Table S5. Number of subjects passing stationarity tests**

**SI Table S6. Parameters of simulated networks**

**SI Table S7. Behavioral scores and descriptions**

**SI Text. Mathematical Note**

